# Evolutionarily Conserved CKI1-Mediated Two-Component Signaling is Required for Female Germline Specification in *Marchantia polymorpha*

**DOI:** 10.1101/2023.09.10.556599

**Authors:** Haonan Bao, Rui Sun, Megumi Iwano, Yoshihiro Yoshitake, Shiori S Aki, Masaaki Umeda, Ryuichi Nishihama, Shohei Yamaoka, Takayuki Kohchi

**Affiliations:** Graduate School of Biostudies, Kyoto University, Kyoto 606-8502, Japan; Graduate School of Science and Technology, Nara Institute of Science and Technology, Ikoma, Nara 630-0192, Japan; Faculty of Science and Technology, Tokyo University of Science, Noda, Chiba 278-8510, Japan

**Keywords:** Female gamete, female germline specification, CKI1, two-component signaling, archegonium development, asymmetric cell division

## Abstract

In land plants, gametes derive from a small number of dedicated haploid cells. ^1^ In angiosperms, one central cell and one egg cell are differentiated in the embryo sac as female gametes for double fertilization, while in non-flowering plants, only one egg cell is generated in the female sexual organ, called the archegonium. ^2,3^ The central cell specification of *Arabidopsis thaliana* is controlled by the histidine kinase CYTOKININ-INDEPENDENT 1 (CKI1), which is a two-component signaling (TCS) activator sharing downstream regulatory components with the cytokinin signaling pathway. ^4–7^ Our phylogenetic analysis suggested that CKI1 orthologs broadly exist in land plants. However, the role of CKI1 in the archegonium-bearing non-flowering plants remains unclear. Here we found that the sole CKI1 ortholog in the liverwort *Marchantia polymorpha*, MpCKI1, which functions through conserved downstream TCS components, regulates the female germline specification for egg cell development in the archegonium. In *M. polymorpha*, the archegonium develops three-dimensionally from a single cell, accumulating MpBONOBO (MpBNB), a master regulator for germline initiation and differentiation. ^8^ We visualized female germline specification by capturing the distribution pattern of MpBNB in the discrete stages of early archegonium development, and found that the MpBNB accumulation is restricted to female germline cells. MpCKI1 is required for the proper MpBNB accumulation in the female germline, and is critical for the asymmetric cell divisions that specify the female germline cells. These results suggest that CKI1-mediated TCS originated during early land plant evolution and participates in female germ cell specification in deeply-diverged plant lineages.

## RESULTS

The histidine kinase CYTOKININ-INDEPENDENT 1 (CKI1) is a master regulator of central cell specification through two-component signaling (TCS) in the embryo sac of *Arabidopsis thaliana*. ^9,10^ To investigate the evolutionary history of CKI1 in archegonium-bearing non-flowering plants, we searched for CKI1 homologs among histidine kinases from publicly available plant genomes and transcriptomes (Table S1), and performed phylogenetic analysis. In addition to those previously reported in seed plants^5,11^, CKI1 orthologs broadly exist in land plant lineages, including ferns, hornworts and liverworts (Figure 1A). In mosses, CKI1 was only found in some basal groups, *e.g. Sphagnum*; in lycophytes, only a few histidine kinases were found in a closely related CKI1-like (CKI1L) clade together with some hornwort sequences; and no CKI1 or CKI1L orthologs were identified in streptophyte algae or other green algae (Figure S1). Our protein alignment highlights the conserved domains in land plant CKI1s, including N-terminal trans-membrane domains, one histidine kinase (HK) domain, one receiver (R) domain, with no apparent cytokinin-receiver CHASE domain (Figure S1). ^12^ Consistent with previous research, the His-Asp (HD) motif is conserved in the HK domain of CKI1, while other histidine kinase families, such as CHK cytokinin receptors and AHK1 osmotic sensors, possess a His-Glu (HE) motif at the same position (Figure S1).^11^

**Figure 1.**
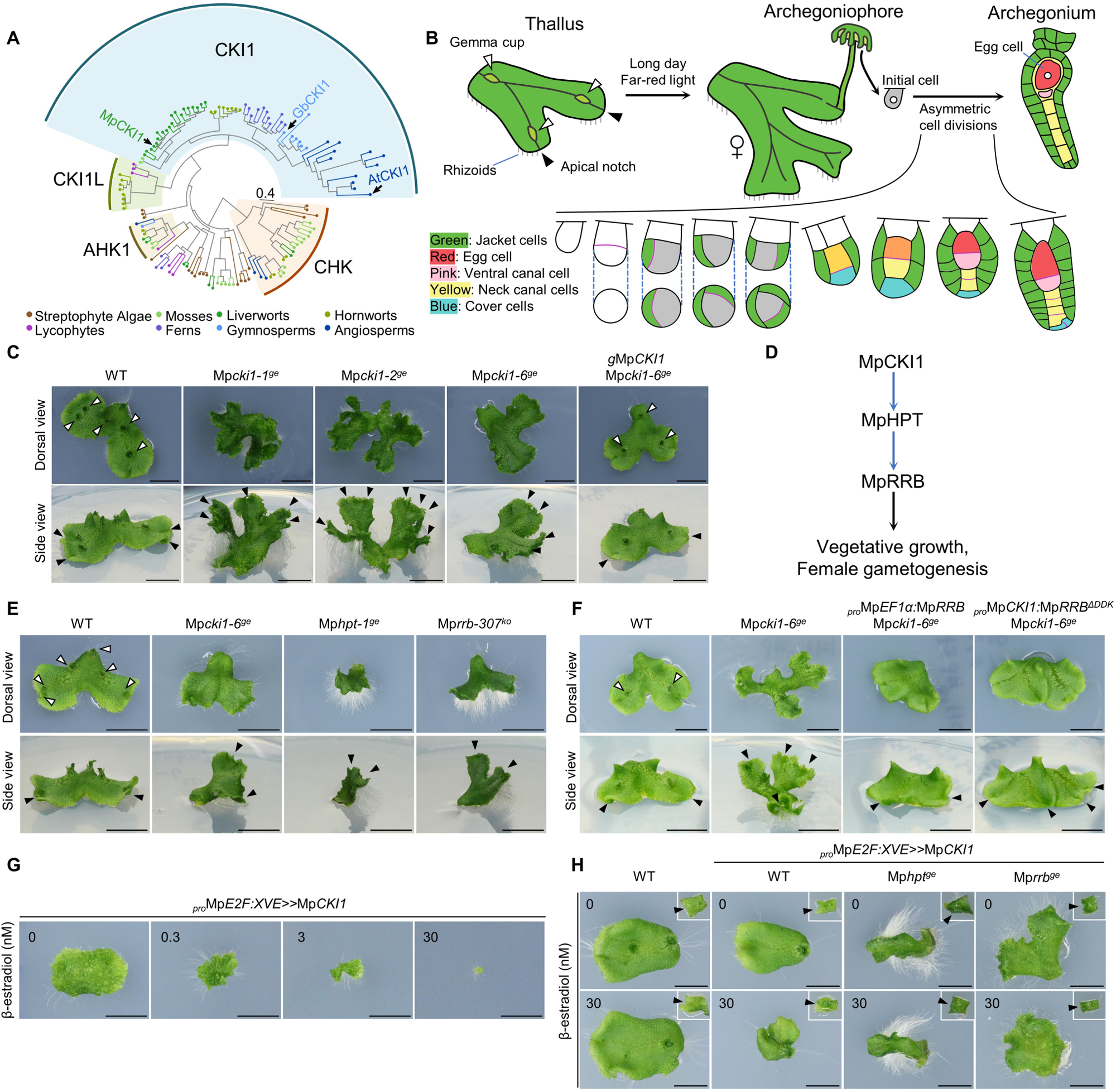
MpCKI1 regulates vegetative growth by conserved downstream TCS components. (A) Phylogenetic tree showing that CKI1 broadly exists in land plants. The scale bar indicates the substitution rate per residue. Mp, *Marchantia polymorpha*; Gb, *Ginkgo biloba*; At, *Arabidopsis thaliana*. AHK1, homologs of ARABIDOPSIS HISTIDINE KINASE1; CHK, CHASE-domain-containing histidine kinase receptors. (B) Schematic diagram of vegetative growth and female sexual reproductive development of *M. polymorpha*. Modified from previous studies. ^3,16,27^ (C) Dorsal and high-angle side view of 14-day-old thalli grown from explants containing apical notches, showing wild-type (WT), several independent Mp*cki1* mutant lines, and a complemented line. (D) Putative MpCKI1-mediated TCS pathway in *M. polymorpha*. Blue arrows indicate phosphorylation, and the black arrow indicates transcriptional regulation. (E) (F) Dorsal and high-angle side views of 14-day-old thalli grown from explants containing apical notches, with genotypes as indicated. (F) Dorsal view of 12-day-old *pro*Mp*E2F:XVE>>*Mp*CKI1* gemmalings grown on medium containing β-estradiol, with concentrations as indicated. Scale bars, 1 mm. (G) Dorsal view of 7-day-old thalli grown from explants with apical notches, with indicated genotypes of inducible MpCKI1 overexpression lines (pro*E2F:XVE*>>Mp*CKI1cds*) in the presence and absence of β-estradiol. Upward bending thalli of Mp*hpt^ge^*and Mp*rrb^ge^* were put down for observation. The origin explants at day 0 are shown with an inset box in the upper right corner on the same scale. Arrowheads indicate apical meristem in the day 0 explants. Scale bars, 10 mm (C, E and F); 5 mm (G and H). Open arrowheads in the dorsal view indicate gemma cups. Arrowheads in the high-angle side view indicate apical notches. See also Figures S1 and S2, as well as Table S1.

In the liverwort *Marchantia polymorpha*, we identified a single CKI1 ortholog (Mp2g02750) and named it MpCKI1. We then generated several independent Mp*cki1* gene-edited mutants (Mp*cki1^ge^*) using the CRISPR-Cas9 system (Figure S2A). ^13–15^ *M. polymorpha* undergoes vegetative growth in a dichotomous branching plant body called thallus, producing gemma cups for asexual reproduction on the dorsal surface and rhizoids from the ventral epidermis (Figure 1B). ^16^ Mp*cki1^ge^* mutants showed abnormal thallus morphology, including upward bending of thallus tips, ectopic serrations on thallus margins, numerous rhizoids, and the absence of gemma cups. These phenotypic features were complemented by introducing a genomic fragment encompassing Mp*CKI1* (*g*Mp*CKI1*) into the Mp*cki1^ge^* mutant (Figures 1C, S2B-S2C). In *A. thaliana*, CKI1 utilizes the downstream regulatory components of cytokinin signaling pathway, including histidine-containing phosphotransfer proteins (HPTs) and type-B response regulators (RRBs), to regulate vegetative growth and female gametogenesis. ^5–7,17^ The *M. polymorpha* genome encodes just one MpHPT and one MpRRB. ^17,18^ We hypothesized that MpCKI1 also functions via these components (Figure 1D). We found that the abnormal thalli of Mp*cki1* mutants resembled Mp*hpt* gene-edited mutants (Mp*hpt^ge^*) generated in this study (Figure S2D), and previously reported Mp*rrb* knock-out (Mp*rrb^ko^*) mutants (Figures 1E, S2E). ^18^ To confirm that MpCKI1 genetically functions upstream of MpRRB, we artificially activated MpRRB signaling in the Mp*cki1^ge^* background by overexpressing Mp*RRB* under a constitutive promoter derived from Mp*ELONGATIONFACTOR1α* (*pro*Mp*EF1α*:Mp*RRB*); alternatively, we used the Mp*CKI1* promoter to express a constitutively activated version of MpRRB with a truncated N-terminal signal receiver domain (*pro*Mp*CKI1*:Mp*RRB^ΔDDK^*). ^19,20^ Both constructs complemented the thallus abnormality in Mp*cki1^ge^* mutant lines except for the gemma cup formation (Figure 1F). In addition, thallus growth was reduced by the overexpression of MpCKI1 with an estrogen-inducible system^21^ (*pro*Mp*E2F:XVE>>*Mp*CKI1*) in a dose-dependent manner (Figure 1G), and the reduction of thallus growth was partially corrected in MpCKI1 overexpressing lines with Mp*hpt^ge^* or Mp*rrb^ge^* mutations (Figure 1H, S2D). Moreover, thallus regeneration was also suppressed by inducible overexpression of MpCKI1, and such suppression was not observed with Mp*hpt^ge^* or Mp*rrb^ge^* lines (Figure S2F). We also performed a transcriptomic analysis with thalli of Mp*cki1^ge^*. This showed a significant overlap of differentially expressed genes with the previously reported *Mprrb^ko^* transcriptome^22^, *e.g.* the gemma cup regulator Mp*GCAM1* (Figure S2G, Table S2). These results suggest that Mp*CKI1* mediates TCS via conserved MpHPT and MpRRB to regulate thallus growth.

To induce sexual reproduction, we grew Mp*cki1* mutants under long-day conditions supplemented with far-red light (FR) irradiation. ^23^ Under such inductive conditions, *M. polymorpha* produces sexual branches called gametangiophores from the apical notch of thalli. In the male branch, called the antheridiophore, spermatogenesis takes place in male sexual organ antheridia inside a disk-like structure. In the female branch, called the archegoniophore, egg-bearing archegonia are formed in the ventral side of the receptacle, at the base of finger-like rays (Figure 1B). ^16^ The male Mp*cki1^ge^* mutants displayed an abnormal morphology of the antheridiophore, resulting in exposed antheridia on the dorsal surface of the antheridiophore, but spermatogenesis was properly completed (Figure S3A). In the female Mp*cki1* mutants, the archegoniophores had shorter and less curved finger-like rays, and only abnormal archegonia-like structures were formed, with no mature egg cells (Figure 2A-2B). The morphology of archegoniophore and egg cell formation were also restored in complementation tests by introducing *g*Mp*CKI1* (Figures 2A-2B). These observations were consistent with the transcriptome profiles of Mp*CKI1*, which showed a notably high expression of Mp*CKI1* in the archegonium (Figure S3B). ^24^ We also performed a transcriptomic analysis of archegoniophores from Mp*cki1* mutants, and we found that several female-enriched genes, including Mp*KNOX1*, Mp*BELL5*, Mp*TRIHELIX28*, Mp*TRIHELIX39*, Mp*AGO4b*, a sexual reproduction related gene, Mp*BZR3*, and an egg cell-specific gene, Mp*EC*, were down-regulated in the archegoniophores of Mp*cki1* mutants (Figure S3C, Table S2). ^25,26^ Taken together, these results suggest that MpCKI1 is essential for archegonium development.

**Figure 2.**
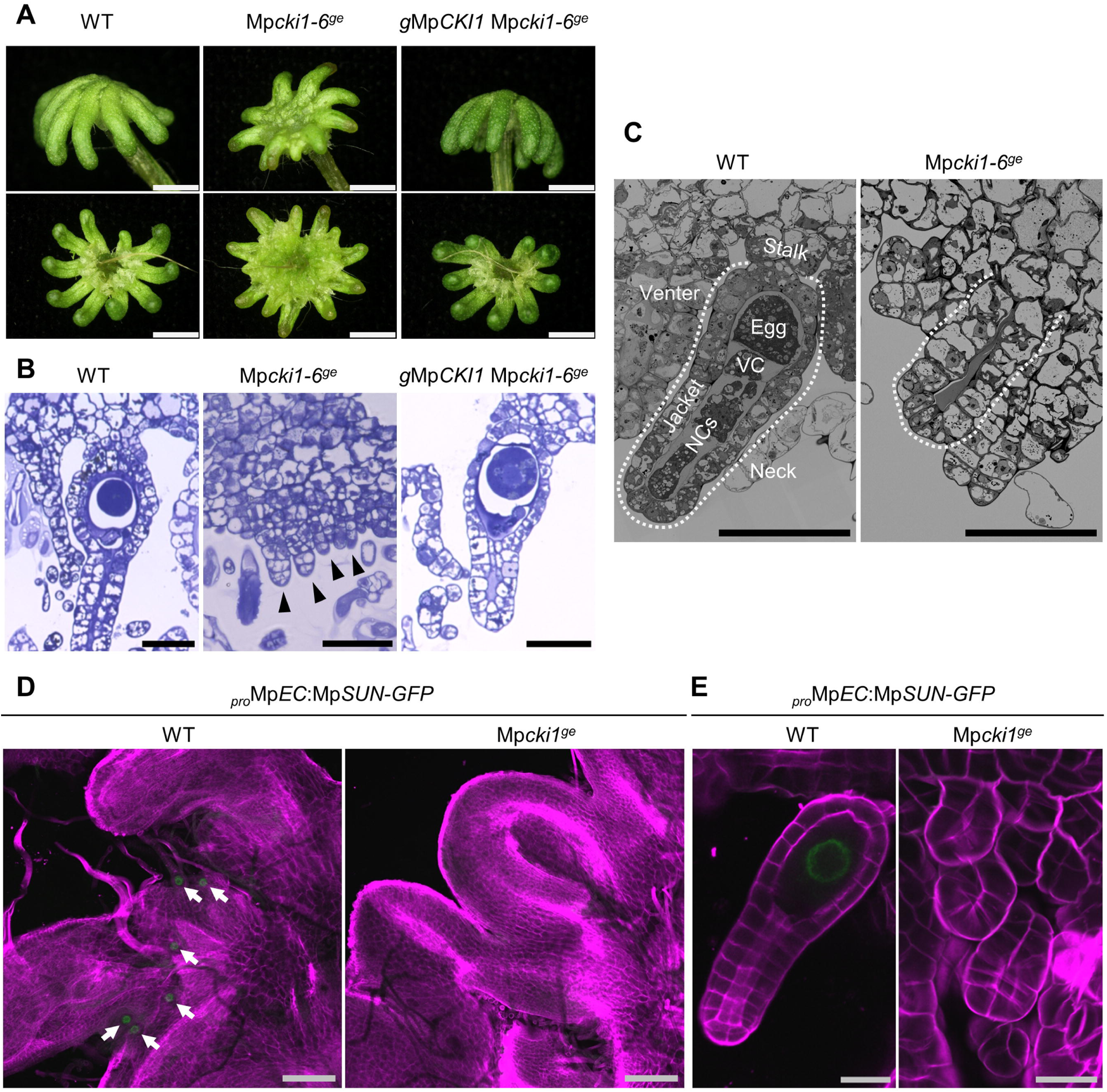
MpCKI1 regulates archegonium development and egg cell differentiation. (A) Side and ventral views of archegoniophores, with genotypes as indicated. Scale bars, 2 mm. (B) Semi-thin microsections of archegoniophores showing archegonia, with the indicated genotypes. Scale bars, 50 μm. Arrowheads indicate abnormal archegonia in Mp*cki1-6^ge^*. (C) Longitudinal sections showing details of archegonium structure, with the indicated genotypes, observed with SEM. Dotted lines indicate outlines of archegonia. Egg, egg cell; VC, ventral canal cell; NCs, neck canal cells. Scale bars, 50 μm. (D) Confocal images showing ventral views of archegoniophores carrying the egg cell marker (*proMpEC*:*MpSUN-GFP*). Arrows indicate egg cell nuclei labeled by GFP (green). Cell walls were stained by Calcofluor White (magenta). Scale bars, 200 μm. (E) Confocal images showing archegonia carrying the egg cell marker. Cell walls were stained by Calcofluor White (magenta). Scale bars, 20 μm. See also Video S1.

To examine the archegonium structure in the Mp*cki1^ge^*mutant, we observed serial longitudinal sections of the archegoniophores with a scanning electron microscope (SEM) (Video S1). In *M. polymorpha*, the archegonium has a long-neck-flask shape, consisting of an egg in its venter, and accessory cells including a ventral canal cell (VC) near the connecting point between the venter and the neck, several neck canal cells (NCs) inside the neck, and cover cells (CVs) at the entrance of the neck (Figure 1B). ^16,27,28^ Developing archegonia with these cell types were observed in wild-type (WT) plants, while in the abnormal archegonia from the Mp*cki1^ge^* mutant, no egg cell, VC, nor NC was formed (Figure 2C). To further confirm that the egg cell was absent in the Mp*cki1* mutants, we disrupted Mp*CKI1* in a reporter line expressing GFP-tagged MpSUN, a nuclear envelope marker, using an egg cell-specific promoter (*pro*Mp*EC*:Mp*SUN-GFP*). ^25^ In the plants carrying the native Mp*CKI1* allele, MpSUN-GFP fluorescence was visible in egg cells at the base of archegoniophore fingers, while in the Mp*cki1^ge^* mutants, only aberrant archegonia lacking MpSUN-GFP fluorescence were observed (Figures 2D-2E). These results suggest that female gametogenesis was defective in Mp*cki1* mutants.

In *M. polymorpha*, the VIIIa basic helix-loop-helix transcription factor MpBONOBO (MpBNB) controls the initiation of gametangia development. During early gametangium development, MpBNB accumulates in the initial cells of gametangia, a subset of their descendants, and finally the immature egg cell. ^8^ To gain insights into the role of Mp*CKI1* in archegonium development, we investigated the influence of Mp*cki1* mutation on the accumulation of MpBNB in Mp*BNB-Citrine* knock-in lines, in which the MpBNB protein is visualized by fusion with the fluorescent protein Citrine. ^8^

The development of archegonia in *M. polymorpha* has been described in previous histological studies, which showed that a single archegonial initial cell undergoes three-dimensional growth with precisely regulated cell divisions in different planes to form a mature archegonium (Figures 1B and S3D). ^3,16,27,28^ We first observed the fluorescence pattern in Mp*BNB-Citrine* expressing archegonia carrying the native Mp*CKI1* allele (Figure 3A). MpBNB-Citrine fluorescence was first detected in the initial cell protruding from the epidermis (Stage I). The initial cell was then partitioned by a transverse wall to produce a two-celled primordium, in which both cells showed MpBNB-Citrine fluorescence (Stage II). The outer cell then underwent three unequal vertical divisions, which separated the cell in the center, with MpBNB-Citrine fluorescence, from the surrounding cells (Stage III-a to c). In the next stage, the MpBNB-Citrine fluorescent cell in the center divided transversely, and produced a proximal cell and a distal cover cell (Stage IV). Following another transverse division of the former proximal cell with MpBNB-Citrine fluorescence, a secondary proximal cell formed, as well as another distal cell in the middle (Stage V). The archegonium then began to differentiate its venter and neck: in the venter, the secondary proximal cell divided unequally into an immature egg cell with MpBNB-Citrine fluorescence and VC; in the neck, MpBNB-Citrine fluorescence was absent, and NCs were formed by successive transverse divisions (Stage VI). Finally, a young archegonium was formed with an egg cell with MpBNB-Citrine fluorescence (Stage VII). In summary, we found that in each asymmetric cell division of female germline cells with accumulated MpBNB during early archegonium development, MpBNB was distributed into two daughter cells, where one of the daughter cells retained the MpBNB accumulation and germline identity, while the other daughter cell eventually lost its MpBNB accumulation and differentiated into the protective structure or accessory cells. Having observed the preferential localization of MpBNB in the egg-cell lineage during female germline specification in the archegonium, we turned our attention to the role of MpCKI1 in this process.

**Figure 3.**
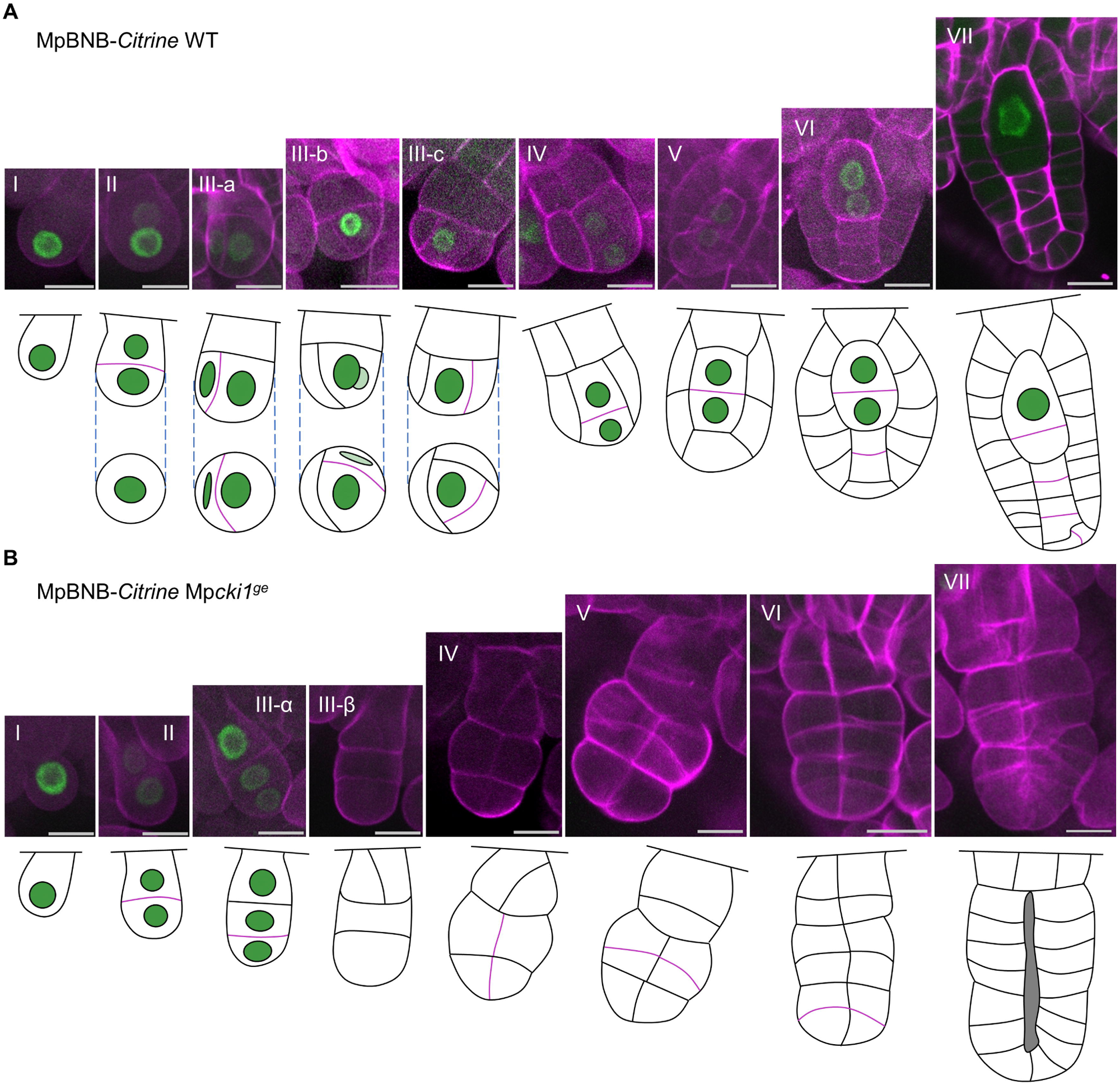
MpCKI1 is required for proper MpBNB accumulation and asymmetric cell divisions in early archegonium development. Confocal images showing MpBNB-Citrine fusion protein distribution in different stages of early archegonium development, obtained from plants carrying the MpCKI1 WT allele (A) or Mp*cki1^ge^* mutants (B). MpBNB accumulation is correlated with Citrine fluorescence (green). Cell walls were stained by Calcofluor White (magenta). Scale bars, 10 μm. Early development of the archegonium is divided into successive stages from (I) to (VII). Interpreted longitudinal and transverse sections of the archegonium are indicated by schematic illustrations. In these illustrations, green circles indicate cell nuclei with relatively intense MpBNB-Citrine signals and purple lines indicate newly formed cell walls in each stage. The growth of the archegonium stalk is omitted. See also Figure S3D.

We monitored the MpBNB-Citrine fluorescence pattern in the archegonium of Mp*cki1^ge^* mutants (Figure 3B). In these plants, the first transverse division of the protruding initial cell occurred normally and produced a two-celled primordium, suggesting that the first cell division of archegonium development was not affected by Mp*cki1^ge^*(Stages I and II). However, the outer cell did not undergo unequal vertical divisions, but instead underwent one transverse division which produced two similar daughter cells, both with MpBNB-Citrine fluorescence; moreover, the basal cell produced in the previous stage still retained MpBNB-Citrine fluorescence (Stage III-α). The MpBNB-Citrine fluorescence failed to be detected in later stages (stage III-β). Subsequently, the cells lacking MpBNB-Citrine fluorescence vertically divided once (Stage IV), then underwent successive transverse divisions (Stages V and VI), and eventually formed an aborted archegonium devoid of MpBNB-Citrine fluorescence (Stage VII). These results suggest that MpCKI1 regulates the female germline specification in two aspects: first, MpCKI1 is required for the asymmetric cell divisions which specify the female germline cells from surrounding protective tissue and accessory cells; second, MpCKI1 restricts the MpBNB accumulation to the female germ cell lineage to establish its identity.

Finally, we tested if MpCKI1 regulates female gametogenesis via TCS. Our observation of Mp*hpt^ge^* and Mp*rrb^ko^* lines showed they produced abnormal archegoniophores with short and less-curved fingers, and formed abnormal archegonia bearing no egg cell (Figure 4A, 4B, and S4A). On the other hand, egg cell formation in Mp*cki1^ge^* was partially restored by artificial activation of MpRRB signaling (Figures 4C, 4D, and S4A). To further characterize the MpCKI1 pathway, several additional mutant lines were evaluated. Putative cytokinin-receptor Mp*chk1^ge^chk2^ge^* double mutants, the cytokinin-activation enzyme Mp*log^ge^* mutants, and transgenic lines overexpressing the cytokinin dehydrogenase Mp*CKX2* produced normal archegoniophores and egg-bearing archegonia, suggesting that CKI1-mediated signaling, but not canonical cytokinin-initiated signaling, is important for female gametogenesis (Figures S4B-S4D). ^17,18^

**Figure 4.**
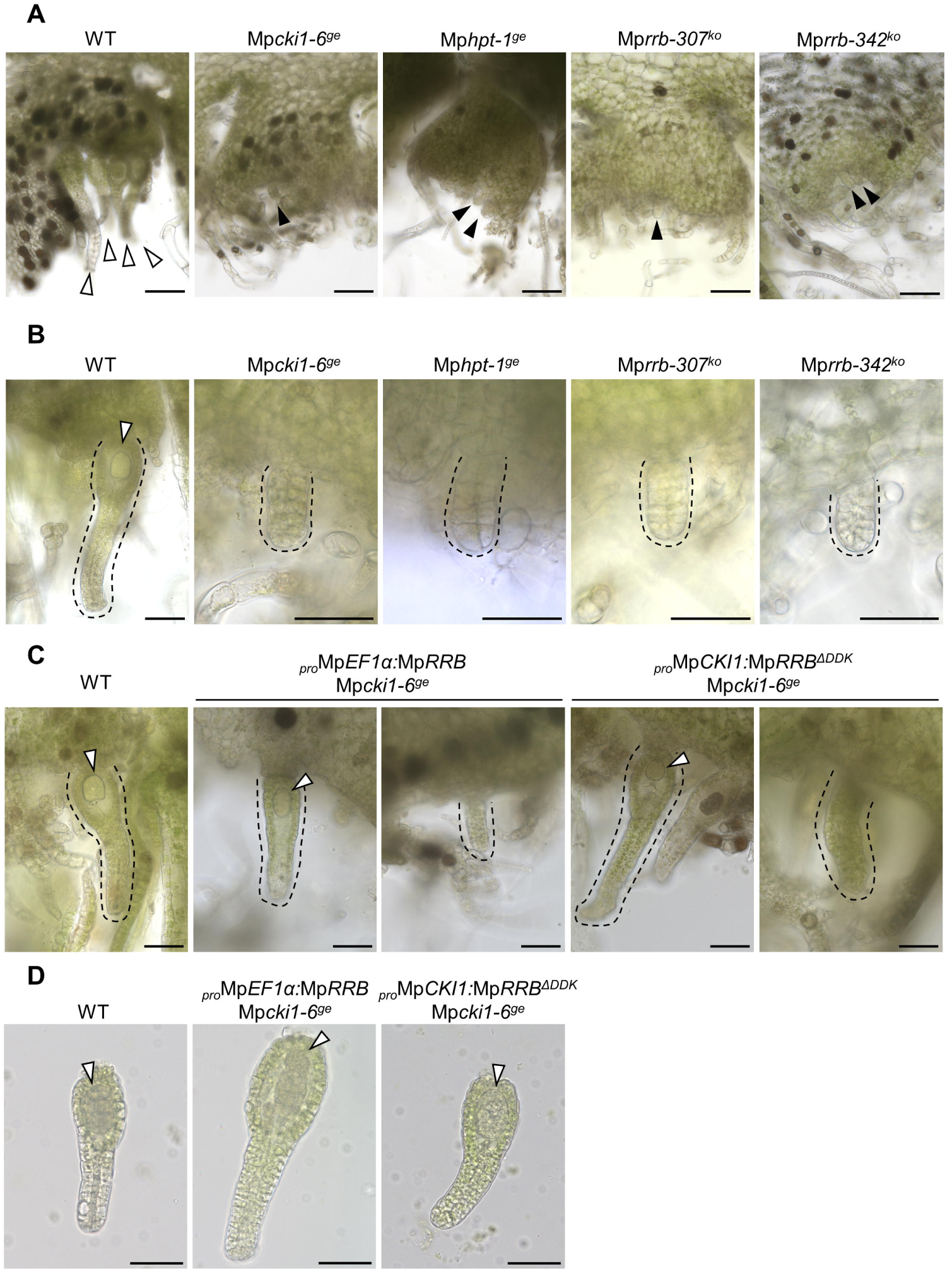
MpCKI1-mediated TCS is essential for female gametogenesis. (A) Archegoniophore sections from plants with the indicated genotypes, showing clusters of archegonia. Scale bars, 100 μm. Open arrowheads indicate archegonia containing an egg cell; arrowheads indicate aborted archegonia without an egg cell. (B) Archegoniophore sections from plants with the indicated genotypes, showing archegonium structure. Scale bars, 50 μm. Black dashed lines indicate outlines of archegonia. Open arrowheads indicate egg cells. (C) Archegoniophore sections from plants with the indicated genotypes, showing partial complementation of Mp*cki1^ge^*by Mp*RRB* gain-of-function. Scale bars, 50 μm. Black dashed lines indicate outlines of archegonia. Open arrowheads indicate the egg cell. (D) Isolated archegonia containing an egg cell, from plants with the indicated genotypes. Scale bars, 50 μm. Open arrowheads indicate egg cells. See also Figure S4.

## DISCUSSION

The germline comprises the cells that are programmed to produce gametes. ^1^ The male germline in angiosperm pollen is specified by asymmetric cell division which produces a vegetative cell and a generative cell. ^1,8^ However, our understanding of the female germline and its specification in the angiosperm embryo sac is limited because of the extremely shortened gametophytic generation with reduced cell number, the coenocytic growth before cellularization, and the differentiation of two dimorphic female gametes. ^1,29^ The angiosperm egg cell fate is determined by positional cues rather than its nuclear lineage, suggesting there is no continuous germline for the angiosperm egg cell, or that the egg cell itself represents a reduced germline. ^1,30^ The other angiosperm female gamete for double fertilization, the central cell, is a defining feature of angiosperms, and plays important roles for their evolutionary success. ^2^ In *A. thaliana*, a master regulator CKI1 regulates the central cell specification: CKI1 is expressed in the megaspore, polarly locates in the chalazal endoplasmic reticulum of the embryo sac at the FG4–FG7 stages, and is distributed to the central cell by nuclear migration and nuclei fusion; furthermore, the ectopic expression of CKI1 in the egg cell precursor can transform it to an additional central cell. ^5,10^

In non-flowering plants, the sole female gamete, the egg cell, is specified by a series of asymmetric cell divisions in the archegonium, which develops from a single initial cell. ^3^ Thus, we may dissect the plant female germline development using a non-flowering model.^1^ Our previous research showed that MpBNB, whose orthologs are essential for male germline specification in *A. thaliana* pollen, functions as a core regulator for both female and male gametangia development in *M. polymorpha.* ^8^ In this study, we visualized *M. polymorpha* female germline specification by the distribution of MpBNB in early archegonium development. Furthermore, we found that MpCKI1 is a key regulator for female germline specification. MpCKI1 restricts the MpBNB accumulation polarly to the female germline cells for the establishment of their identity. Particularly, in stage III of archegonium development, we observed that the orientation of the cell division plane for the asymmetric cell division was changed in Mp*cki1* mutants, suggesting that Mp*cki1* mutants failed to establish the polarity for proper archegonium development. In the embryo sac of *A. thaliana*, polarly localized CKI1 is essential for the establishment or maintenance of a micropylar-chalazal polarity. ^5^ Taken together, CKI1 may participate in a common regulation system for polarity in female germ cell specification of land plants.

Interestingly, female genetic transmission in *A. thaliana* was not significantly affected by mutations in *BNB* orthologs, indicating a different or additional mechanism for the female germline specification in angiosperms. ^8^ Indeed, the angiosperm egg apparatus is derived from nuclei that do not express CKI1 in FG4, and *cki1* mutated embryo sacs are still able to produce fertile egg cells, indicating that egg cell differentiation is independent of CKI1 in angiosperm embryo sac development. ^5,10^ In *M. polymorpha*, by contrast, MpCKI1 is required for the egg cell formation, and the activation of MpCKI1-mediated TCS restored egg cell formation in Mp*cki1* mutants. These differences suggest that CKI1-mediated TCS is involved in the specification of different female gametes in the embryo sac of *A. thaliana* and the archegonium of *M. polymorpha*.

Our phylogenetic analysis suggests that CKI1 is broadly present in land plant lineages. As the downstream TCS components predated the emergence of CKI1 and are conserved in land plants^31^, CKI1-mediated TCS could be a regulatory system broadly used for female gametogenesis in land plants. The origin of CKI1 in the common ancestor of land plants may play an important role in the innovation of the archegonium, a multicellular sexual organ with three-dimensional growth to protect and support female gametogenesis, fertilization, and embryogenesis in terrestrial environments. ^32^ Moreover, the switch of MpCKI1’s role between egg cell and central cell specification may coincide with the origin of angiosperms. Further studies on the function of CKI1 in other land plant lineages may shed light on these key evolutionary events, and understanding how CKI1 regulates cell polarity may provide insights into the genetic bases of female germline specification in the archegonium and embryo sac development.

## Acknowledgements

We thank Tatsuya Katsuno for technical assistance with SEM; Tetsuya Hisanaga for sharing the egg cell marker line (*pro*Mp*EC*:MpSUN*-GFP*); Yukari Sando and Takefumi Kondo for NGS and RNA sequencing; and James Alan Hejna for critically reading the manuscript. This work was supported by MEXT KAKENHI grant number JP19H05675 (T.K.), JP19H04860, JP20H05780 (S.Y), and JSPS KAKENHI grant numbers JP17H07424, JP22H00417, JP22K19345 (T.K.).

## Author contributions

Conceptualization, H.B., S.Y. and T.K.; Methodology, H.B., R.S., M.I., Y.Y., S.Y., R.N., and T.K.; Validation, S.Y. and T.K.; Investigation, H.B., R.S., M.I., and Y.Y.; Resources, S.S.A. and M.U.; Writing – Original draft, H.B.; Writing – Reviewing and Editing, H.B., R.S., Y.Y., S.S.A., R.N. S.Y. and T.K.; Funding Acquisition, S.Y. and T.K.

## METHODS

### Lead contact

Requests for further information, or for resources and reagents, should be directed to and will be fulfilled by the lead contact, Takayuki Kohchi.

### Materials availability

Plasmids and plant materials generated in this research are all available from the lead contact upon request. Please note that the distribution of transgenic plants will be governed by material transfer agreements (MTAs) and will be dependent on appropriate import permits acquired by the receiver.

### Data and code availability

Genome and RNA-seq data have been deposited in the Sequence Read Archive at DNA Data Bank of Japan (DDBJ) under Bioproject DRA016515, DRA016516 and are publicly available as of the date of publication. Original code used for phylogenetic analysis and RNA-seq analysis was deposited at https://github.com/dorrenasun/MpCKI1 and is publicly available as of the date of publication. Any additional information required to reanalyze the data reported in this research is available from the lead contact upon request.

## EXPERIMENTAL MODEL AND SUBJECT DETAILS

### *M. polymorpha* lines and growth conditions

The female accession Takaragaike-2 (Tak-2) and the male Takaragaike-1 (Tak-1) of *Marchantia polymorpha subsp. ruderalis* were used in this study as wild-type plants. ^33^ Plants were grown on half-strength Gamborg’s B5 medium^34^ containing 1% (w/v) agar or 0.6% (w/v) gellan gum (for rhizoid observation) under continuous white light (50-60 mmol photons m^-2^ s^-1^; CCFL OPT-40C-N-L, Optorom) at 22°C. For estrogen-inducible MpCKI1 overexpression, gemmae or thallus cut with/without an apical notch were grown on half-strength Gamborg’s B5 medium containing 0.03 nM to 30 nM β-estradiol. To induce reproductive growth, 10-day-old gemmalings or thallus cuts (from the lines which cannot produce gemma) were transferred to half-strength Gamborg’s B5 medium plus 1% (w/v) sucrose under long-day (16 hours in light and 8 hours in dark) conditions supplemented with far-red light (20-30 mmol photons m^-2^ s^-1^, VBL-TFL600-IR730, Valore, Kyoto, Japan).

## METHOD DETAILS

### Phylogenetic analysis and protein domain prediction

To find CKI1 homologs in Viridiplantae, CKI1s from *A. thaliana* (AT2G47430) and *G. biloba* (chr3.1901) were queried with protein-protein BLAST (ver. 2.6.0) ^35,36^ against sequences from publicly available genomes (see Table S1 for a full list of sources and references), as well as selected transcriptomes from the 1000 plant transcriptomes initiative (1KP) and a previous publication. ^37–39^ All hits with e-values lower than 0.001 were retrieved and analyzed with HMMER (ver. 3.3.2) to identify HK (PF00512 and PF02518) and receiver domains (PF00072) (“HMMER Webpage” 2023). Sequences that contained HK domains were aligned with MAFFT (ver. 7.520; algorithm: FFT-NS-2). ^40^ After masking positions with >80% gaps, an initial tree covering a broad range of HKs was inferred from the multiple sequence alignment using IQ-TREE (ver. 2.0.3) with automatic model selection (ModelFinder; LG+G4 chosen according to the Bayesian Information Criterion) and 1000 SH-aLRT branch tests. ^41–43^ From this initial tree, candidate CKI1 homologs and sequences from closely-related clades were selected for further analysis. A new multiple sequence alignment was built with MAFFT (ver. 7.520; algorithm: E+INSI) and masked for >80% gap positions. ^40^ The final phylogenetic inference was carried out with IQ-TREE with automatic model selection (ModelFinder; JTT+F+I+G4 chosen) and 1000 standard nonparametric bootstraps. ^41,42^ CHKs from representative genomes were used as the outgroup to root the tree. For a complete search of protein domains, sequences in the final tree were analyzed with InterProScan (ver. 5.61-93.0). ^44,45^ The phylogenetic tree and domain organizations were visualized using the ggtree R package suite (ver. 3.6.2). ^46–48^ Overlapping, redundant domain annotations were merged for a concise presentation. The data sources, references, and notes for sequence selection are listed in Table S1. The scripts and key output files are available at https://github.com/dorrenasun/MpCKI1.

### Plasmid construction for genome editing and complementation tests

Gateway vectors with selection markers, reporters and tags used in this study were described previously. ^49,50^ To generate Mp*cki1^ge^*, Mp*hpt^ge^*, Mp*rrb^ge^*, Mplog*^ge^* and Mp*chk1^ge^chk2^ge^* mutants, single guide RNA (sgRNA) sequences were inserted into pMpGE_En03, and were then subcloned into the binary vector pMpGE010 or pMpGE011 using Gateway LR Clonase II Enzyme mix (Thermo Fisher Scientific). For Mp*cki1-6^ge^*, two pairs of sgRNAs were inserted separately into pMpGE_En04, pBC-GE12, pBC-GE23, and pBC-GE34, then fragments from these vectors were combined to create an entry clone with cassettes for all four sgRNAs. ^14,15^ The sgRNA cassettes were then transferred into pMpGE018 by the Gateway LR reaction.

The promoter region, coding sequence (CDS), and the genomic fragment (including promoter and coding region) of Mp*CKI1* were amplified from Tak-1 genomic DNA by PCR using KOD One PCR Master Mix (Toyobo Life Science) and the following primer sets: MpCKI1_pro_FB4 and MpCKI1_pro_RB1R (promoter region); MpCKI1-CDS-F and MpCKI1-CDS-R (CDS); MpCKI1_5’_EcoRI_IFF and MpCKI1_3’_EcoRI_IFR (genomic region). The promoter region was inserted into pDONAp4-p1R using Gateway BP Clonase II Enzyme mix (Thermo Fisher Scientific), whereas the CDS was directionally inserted into pENTR/D-TOPO and the genomic fragment was inserted into the pENTR1A vector (Thermo Fisher Scientific) by conventional cloning methods. For Mp*cki1-6^ge^* complementation, the genomic fragment in pENTR1A was transferred into pMpGWB101 by a Gateway LR reaction. For MpCKI1 inducible overexpression (*pro*Mp*E2F:XVE*>>Mp*CKI1cds*), the Mp*CKI1* CDS in pENTR/D-TOPO was recombined with pMpGWB168.

Mp*RRB^ΔDDK^* coding sequences (CDSs) were amplified by PCR from Tak-1 thallus complementary DNA (cDNA) using primer set MpRRBΔDDK-CACCATG-F and MpRRB-nstop-R (Mp*RRB^ΔDDK^cds*). First, the cDNA was prepared by RNA extraction using an RNeasy Plant Mini kit (Qiangen) and reverse transcription using ReverTra Ace (Toyobo Life Science). After amplification, the PCR products were directionally cloned into the pENTR/D-TOPO vector (Thermo Fisher Scientific) to create entry vectors. For Mp*cki1-6^ge^* complementation by MpRRB overexpression, the Mp*RRBcds* cassette was transferred into pMpGWB103 using a Gateway LR reaction. For Mp*cki1-6^ge^* complementation by Mp*RRB^ΔDDK^*, the MpCKI1pro cassette in pDONAp4-p1R and the Mp*RRB^ΔDDK^cds* cassette in pENTR/D-TOPO were transferred into R4pMpGWB139 by a Gateway LR reaction.

### Plant transformation

Sporelings or regenerating thalli of *M. polymorpha* were transformed with binary vectors using *Agrobacterium tumefaciens* GV2260 following previously described methods. ^51,52^ Young thallus fragments (from lines that cannot produce gemmae) or 12-day-old gemmalings were picked up and cut into small pieces without apical notches, and cultured for 3 days on half-strength Gamborg’s B5 medium with 1% (w/v) agar and 1% (w/v) sucrose for regeneration. *Agrobacterium* carrying the relevant binary vector was then co-cultivated with regenerating thalli in M51C medium containing 100 mM acetosyringone for 3 days, with agitation. ^53^ Co-cultivated thalli were then washed with sterile water and placed on selective media with 100 μg mL^-1^ cefotaxime to kill remaining agrobacterium and 10 μg mL^-1^ hygromycin or 0.5 μM chlorsulfuron, depending on the selection markers in the binary vectors.

### Genome sequencing of Mp*cki1-6ge*

Genomic DNA was extracted from thalli of Mp*cki1-6^ge^*by a previously described polyvinylpolypyrrolidone (PVPP) method, with slight modifications. ^54^ Plant tissue was frozen in liquid nitrogen and crushed into powder, then incubated in 2mL of PVPP buffer (50 mM Tris-HCl [pH9.5], 10 mM EDTA, 4 M NaCl, 1% (w/v) CTAB, 0.5% (w/v) PVPP, 1% (v/v) 2-mercaptoethanol) at 80 °C for 30 min in a water bath. After vortexing with 3 mL of chloroform and 2 mL of TE-saturated phenol, the samples were centrifuged at 11,000 × g for 5 minutes at room temperature. The upper (aqueous) phase was transferred to a new tube, 3 mL of sterile water and 6 mL of 100% ethanol were then added, and the tube was placed at -80°C until frozen. After 2 hours, the samples were thawed, centrifuged at 20,000 × g for 15 minutes at 4 °C, and the supernatant was discarded. The pellets were dissolved in 2 mL of TE buffer and heated at 60 °C for 10 minutes. To dissolve the DNA from the pellets, 2 mL TE buffer was added and heated at 60°C for 10 minutes. The TE buffer with dissolved DNA was then carefully collected without disturbing the remaining pellets, and treated with 2 μL of RNaseA (10 mg mL^-1^) for 5 minutes at 37°C. The DNA was then purified by ethanol precipitation. Genomic DNA libraries were prepared using an NEBNext Ultra II FS DNA Library Prep Kit for Illumina (New England Biolabs) according to the manufacturer’s instructions. The quality of DNA was evaluated by an Agilent 2100 Bioanalyzer system and Qubit fluorometer (Invitrogen). All genomic DNA libraries were sequenced as 150-nucleotide paired-end reads using an Illumina NextSeq 550 system. The sequence reads were mapped to the *M. polymorpha* genome (v5.1) by Bowtie2 and inspected in the Integrative Genomics Viewer. ^55–57^

### Histology and Microscopy

To prepare semi-thin and thin sections, mature archegoniophores and antheridiophores with elongated stalks were fixed with 2.5% (v/v) glutaraldehyde (GA) and 2% (w/v) paraformaldehyde (PFA) in 50 mM phosphate buffer (PB, pH 7.2) at 4℃ for more than 3 hours. After washing with PB, the samples were post-fixed with 2% (w/v) osmium tetroxide at room temperature for 2 hours, washed 3 times with an 8% (w/v) sucrose solution for 30 minutes each, dehydrated in a graded series of ethanol concentrations, and embedded in epoxy resin (Quetol 812; Nisshin EM, Tokyo, Japan). ^14^ For light microscopy, semi-thin sections were cut using an EM-UC6 ultramicrotome (Leica, Wetzlar, Germany), mounted on microscope slides, and stained with 1% (w/v) toluidine blue. Samples were observed under bright field with a BX-700 microscope (Keyence, Osaka, japan). For SEM observation, serial sections (250-nm thick) were cut with a diamond knife using an EM-UC6 ultramicrotome, mounted on a silicon wafer (380-µm thick; Canosis, Tokyo, Japan) and stained with 2% (w/v) uranyl acetate and a lead stain solution. Samples were observed with a JSM-7900F SEM (JEOL, Tokyo, Japan). Images were obtained at 4 kV. Stacked images were aligned using the Stacker NEO software (JEOL, Tokyo, Japan).

For the morphological observation of gametangiophores, samples were photographed with an Olympus SZX16 stereomicroscope. For the SEM of thallus margins, thallus fragments of 3-week-old plants were flash-frozen in liquid nitrogen and observed with a HITACHI Miniscope TM3000.

For sperm observation, antheridia were picked up, put on a slide glass and disrupted in water to release the sperm. The sperm were observed with a BX-700 microscope (Keyence, Osaka, japan).

To observe archegonium clusters with agar sectioning, archegoniophores were embedded in 5% (w/v) agar, sectioned with a vibratome (DOSAKA LinearSlicer Pro7), and observed with a BX-700 microscope (Keyence, Osaka, japan).

### Fluorescence microscopy

For egg cell marker observation, archegoniophores in the tips of elongated stalks were used. For archegonium development of WT and Mp*cki1* lines, developing archegoniophores without stalk elongation were used. Archegoniophores were fixed in 4% (w/v) PFA in 1 x phosphate-buffered saline (PBS, pH 7.4), treated with ClearSeeAlpha solution (ClearSee solution containing 50 mM sodium sulfite) overnight, then stained with ClearSee solution containing 0.2 mg/mL Calcofluor White for over 3 hours at room temperature. Images were obtained with a FLUOVIEW FV1000 confocal laser scanning microscope (CLSM; Olympus, Tokyo, Japan). ^58,59^ For GFP fluorescence, the excitation wavelength was 488 nm and emission wavelengths 500-525 nm; for Citrine fluorescence, the excitation wavelength was 515 nm and emission wavelengths 530-570 nm; for Calcofluor-white-stained cell walls, the excitation wavelength was 405 nm and emission wavelengths 425-475 nm.

### RNA sequencing

Total RNA was extracted from two-week-old thallus tips and young archegoniophores without elongated stalks of Tak-2 and Mp*cki1-6^ge^*lines by the RNeasy Plant Mini kit (QIAGEN). For each material, three samples from different plants were collected as biological replicates. The quality was checked by an Agilent 2100 Bioanalyzer (Agilent) with an RNA 6000 nano kit, and concentrations were determined with a Qubit fluorometer (Invitrogen). The mRNA was enriched with an NEBNext Poly(A) mRNA Magnetic Isolation Module (New England Biolabs). The libraries were prepared with an NEBNext Ultra II Directional RNA Library Prep Kit for Illumina (New England Biolabs) and amplified with an NEBNext Multiplex Oligos kit for Illumina (New England Biolabs) according to the manufacturer’s instructions. All transcriptome libraries were sequenced as 75-nucleotide single reads using an Illumina NextSeq 550 system.

For RNA-seq data processing, reads were mapped to the *M. polymorpha* reference genome (MpTak_v6.1) with the V-chromosome removed, and quantified by Salmon v1.9.0. ^60,61^ The R package DESeq2 (version 1.38.3) was used for differential gene expression analysis. ^62^ In pairwise analyses, genes with a minimum fold change of 1.5 and an adjusted *p*-value lower than 0.01 (Wald test with Benjamini-Hochberg adjustment) were considered as significantly changed in expression. For the comparison of thallus transcriptome profiles, RNA-seq data from Mp*rrb-307^ko^* and WT plants, collected in previous research, were retrieved and re-analyzed following the same pipeline above. ^22^ Fisher’s exact test was used to test if the overlap between two gene sets was significant.

## QUANTIFICATION AND STATISTICAL ANALYSIS

For imaging analyses, at least three individuals representing at least two independent transgenic lines were evaluated, otherwise the details are described in the figure legends.

