## Supplementary material for "Evolutionarily Conserved CKI1-Mediated Two-Component Signaling is Required for Female Germline Specification in *Marchantia polymorpha*": Figure S1-S4, Table S3

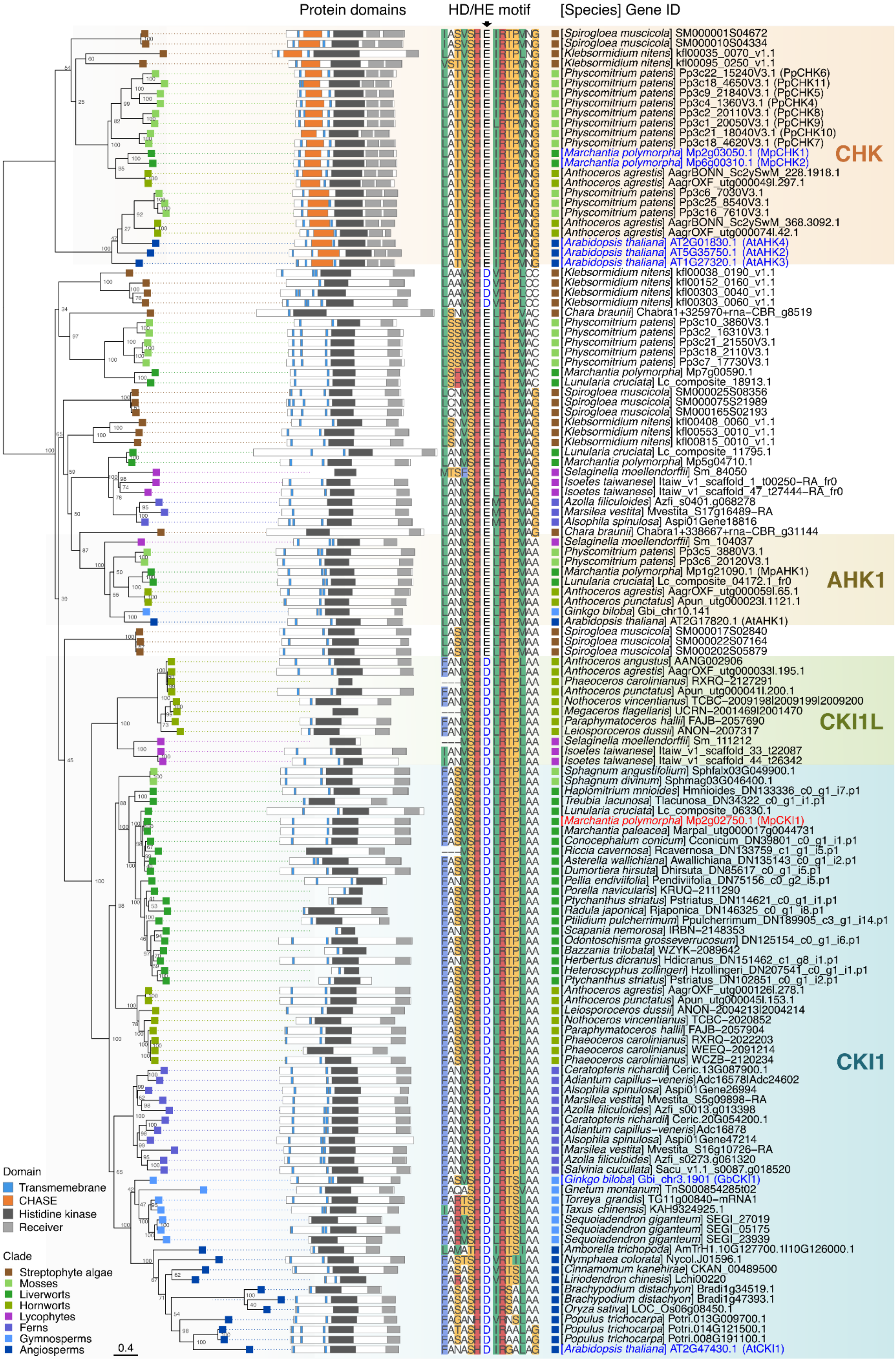

Protein domains
HD/HE motif
[Species] Gene ID
[Spirogloea muscicola] SM000001S04672
[Spirogloea muscicola] SM000010S04334
[Klebsormidium nitens] kfl00035_0070_v1.1
[Klebsormidium nitens] kfl00095_0250_v1.1
[Physcomitrium patens] Pp3c22_15240V3.1 (PpCHK6)
[Physcomitrium patens] Pp3c18_4650V3.1 (PpCHK11)
[Physcomitrium patens] Pp3c9_21840V3.1 (PpCHK5)
[Physcomitrium patens] Pp3c4_1360V3.1 (PpCHK4)
[Physcomitrium patens] Pp3c2_20110V3.1 (PpCHK8)
[Physcomitrium patens] Pp3c1_20050V3.1 (PpCHK9)
[Physcomitrium patens] Pp3c21_18040V3.1 (PpCHK10)
[Physcomitrium patens] Pp3c18_4620V3.1 (PpCHK7)
[Marchantia polymorpha] Mp2g03050.1 (MpCHK1)
[Marchantia polymorpha] Mp6g00310.1 (MpCHK2)
[Anthoceros agrestis] AagrBONN_Sc2ySwM_228.1918.1
[Anthoceros agrestis] AagrOXF_utg000049l.297.1
[Physcomitrium patens] Pp3c6_7030V3.1
[Physcomitrium patens] Pp3c25_8540V3.1
[Physcomitrium patens] Pp3c16_7610V3.1
[Anthoceros agrestis] AagrBONN_Sc2ySwM_368.3092.1
[Anthoceros agrestis] AagrOXF_utg000074l.42.1
[Arabidopsis thaliana] AT2G01830.1 (AtAHK4)
[Arabidopsis thaliana] AT5G35750.1 (AtAHK2)
[Arabidopsis thaliana] AT1G27320.1 (AtAHK3)
CHK
[Klebsormidium nitens] kfl00038_0190_v1.1
[Klebsormidium nitens] kfl00152_0160_v1.1
[Klebsormidium nitens] kfl00303_0040_v1.1
[Klebsormidium nitens] kfl00303_0060_v1.1
[Chara braunii] Chabra1+325970+rna-CBR_g8519
[Physcomitrium patens] Pp3c10_3860V3.1
[Physcomitrium patens] Pp3c2_16310V3.1
[Physcomitrium patens] Pp3c21_21550V3.1
[Physcomitrium patens] Pp3c18_2110V3.1
[Physcomitrium patens] Pp3c7_17730V3.1
[Marchantia polymorpha] Mp7g00590.1
[Lunularia cruciata] Lc_composite_18913.1
[Spirogloea muscicola] SM000025S08356
[Spirogloea muscicola] SM000075S21989
[Spirogloea muscicola] SM000165S02193
[Klebsormidium nitens] kfl00408_0060_v1.1
[Klebsormidium nitens] kfl00553_0010_v1.1
[Klebsormidium nitens] kfl00815_0010_v1.1
[Lunularia cruciata] Lc_composite_11795.1
[Marchantia polymorpha] Mp5g04710.1
[Selaginella moellendorffii] Sm_84050
[Isoetes taiwanese] Itaiw_v1_scaffold_1_t00250-RA_fr0
[Isoetes taiwanese] Itaiw_v1_scaffold_47_t27444-RA_fr0
[Azolla filiculoides] Azfi_s0401.g068278
[Marsilea vestita] Mvestita_S17g16489-RA
[Alsophila spinulosa] Aspi01Gene18816
[Chara braunii] Chabra1+338667+rna-CBR_g31144
[Selaginella moellendorffii] Sm_104037
[Physcomitrium patens] Pp3c5_3880V3.1
[Physcomitrium patens] Pp3c6_20120V3.1
[Marchantia polymorpha] Mp1g21090.1 (MpAHK1)
[Lunularia cruciata] Lc_composite_04172.1_fr0
[Anthoceros agrestis] AagrOXF_utg000059l.65.1
[Anthoceros punctatus] Apun_utg000023l.1121.1
[Ginkgo biloba] Gbi_chr10.141
[Arabidopsis thaliana] AT2G17820.1 (AtAHK1)
AHK1
[Spirogloea muscicola] SM000017S02840
[Spirogloea muscicola] SM000022S07164
[Spirogloea muscicola] SM000202S05879
[Anthoceros angustus] AANG002906
[Anthoceros agrestis] AagrOXF_utg000033l.195.1
[Phaeoceros carolinianus] RXRQ-2127291
[Anthoceros punctatus] Apun_utg000041l.200.1
[Nothoceros vincentianus] TCBC-2009198|2009199|2009200
[Megaceros flagellaris] UCRN-2001469|2001470
[Paraphymatoceros hallii] FAJB-2057690
[Leiosporoceros dussii] ANON-2007317
[Selaginella moellendorffii] Sm_111212
[Isoetes taiwanese] Itaiw_v1_scaffold_33_t22087
[Isoetes taiwanese] Itaiw_v1_scaffold_44_t26342
CKI1L
[Sphagnum angustifolium] Sphfalx03G049900.1
[Sphagnum divinum] Sphmag03G046400.1
[Haplomitrium mnioides] Hmnioides_DN133336_c0_g1_i7.p1
[Treubia lacunosa] Tlacunosa_DN34322_c0_g1_i1.p1
[Lunularia cruciata] Lc_composite_06330.1
[Marchantia polymorpha] Mp2g02750.1 (MpCKI1)
[Marchantia paleacea] Marpal_utg000017g0044731
[Conocephalum conicum] Cconicum_DN39801_c0_g1_i1.p1
[Riccia cavernosa] Rcavernosa_DN133759_c1_g1_i5.p1
[Asterella wallichiana] Awallichiana_DN135143_c0_g1_i2.p1
[Dumortiera hirsuta] Dhirsuta_DN85617_c0_g1_i5.p1
[Pellia endiviifolia] Pendiviifolia_DN75156_c0_g2_i5.p1
[Porella navicularis] KRUQ-2111290
[Ptychanthus striatus] Pstriatus_DN114621_c0_g1_i1.p1
[Radula japonica] Rjaponica_DN146325_c0_g1_i8.p1
[Ptilidium pulcherrimum] Ppulcherrimum_DN189905_c3_g1_i14.p1
[Scapania nemorosa] IRBN-2148353
[Odontoschisma grosseverrucosum] DN125154_c0_g1_i6.p1
[Bazzania trilobata] WZYK-2089642
[Herbertus dicranus] Hdicranus_DN151462_c1_g8_i1.p1
[Heteroscyphus zollingeri] Hzollingeri_DN207541_c0_g1_i1.p1
[Ptychanthus striatus] Pstriatus_DN102851_c0_g1_i2.p1
[Anthoceros agrestis] AagrOXF_utg000126l.278.1
[Anthoceros punctatus] Apun_utg000045l.153.1
[Leiosporoceros dussii] ANON-2004213|2004214
[Nothoceros vincentianus] TCBC-2020852
[Paraphymatoceros hallii] FAJB-2057904
[Phaeoceros carolinianus] RXRQ-2022203
[Phaeoceros carolinianus] WEEQ-2091214
[Phaeoceros carolinianus] WCZB-2120234
[Ceratopteris richardii] Ceric.13G087900.1
[Adiantum capillus-veneris]Adc16578|Adc24602
[Alsophila spinulosa] Aspi01Gene26994
[Marsilea vestita] Mvestita_S5g09898-RA
[Azolla filiculoides] Azfi_s0013.g013398
[Ceratopteris richardii] Ceric.20G054200.1
[Adiantum capillus-veneris]Adc16878
[Alsophila spinulosa] Aspi01Gene47214
[Marsilea vestita] Mvestita_S16g10726-RA
[Azolla filiculoides] Azfi_s0273.g061320
[Salvinia cucullata] Sacu_v1.1_s0087.g018520
[Ginkgo biloba] Gbi_chr3.1901 (GbCKI1)
[Gnetum montanum] TnS000854285t02
[Torreya grandis] TG11g00840-mRNA1
[Taxus chinensis] KAH9324925.1
[Sequoiadendron giganteum] SEGI_27019
[Sequoiadendron giganteum] SEGI_05175
[Sequoiadendron giganteum] SEGI_23939
[Amborella trichopoda] AmTrH1.10G127700.1I10G126000.1
[Nymphaea colorata] Nycol.J01596.1
[Cinnamomum kanehirae] CKAN_00489500
[Liriodendron chinesis] Lchi00220
[Brachypodium distachyon] Bradi1g34519.1
[Brachypodium distachyon] Bradi1g47393.1
[Oryza sativa] LOC_Os06g08450.1
[Populus trichocarpa] Potri.013G009700.1
[Populus trichocarpa] Potri.014G121500.1
[Populus trichocarpa] Potri.008G191100.1
[Arabidopsis thaliana] AT2G47430.1 (AtCKI1)
CKI1
Domain
Transmembrane
CHASE
Histidine kinase
Receiver
Clade
Streptophyte algae
Mosses
Liverworts
Hornworts
Lycophytes
Ferns
Gymnosperms
Angiosperms
0.4

**Figure S1. Phylogenetic analysis of CKI1 and other histidine kinases (related to Figure. 1)**

To infer this phylogenetic tree as described in the STAR Methods section, a subset of protein sequences was selected from the initial analysis to include the ancestral node of all CKI1 homologs and closely-related *M. polymorpha* proteins, as well as CHASE-containing histidine kinase receptors (CHKs, as the outgroup). Numbers next to internal nodes represent percentage support values from 1000 standard non-parametric bootstraps by IQ-TREE 2. The length of each branch is related to the number of substitutions per residue, as indicated by the scale bar. The schematic diagrams in the middle left show the domain structures of each peptide sequence, which is predicted by InterProScan. The core sequences from the histidine kinase (HK) domain are aligned in the middle right. This motif is highly conserved among CKI1 homologs from non-seed plants, but is more variable in those from seed plants. All the sequences from CKI1 and CKI1L clades share a conserved aspartate residue (D) next to the histidine phosphorylation site. Homologs of CKI1, CKI1-like (CKI1L), CHK and ARABIDOPSIS HISTIDINE KINASE1 (AHK1) are shaded with different colors. The sequence name of MpCKI1 is highlighted with red, and several sequences of interest are highlighted with blue.

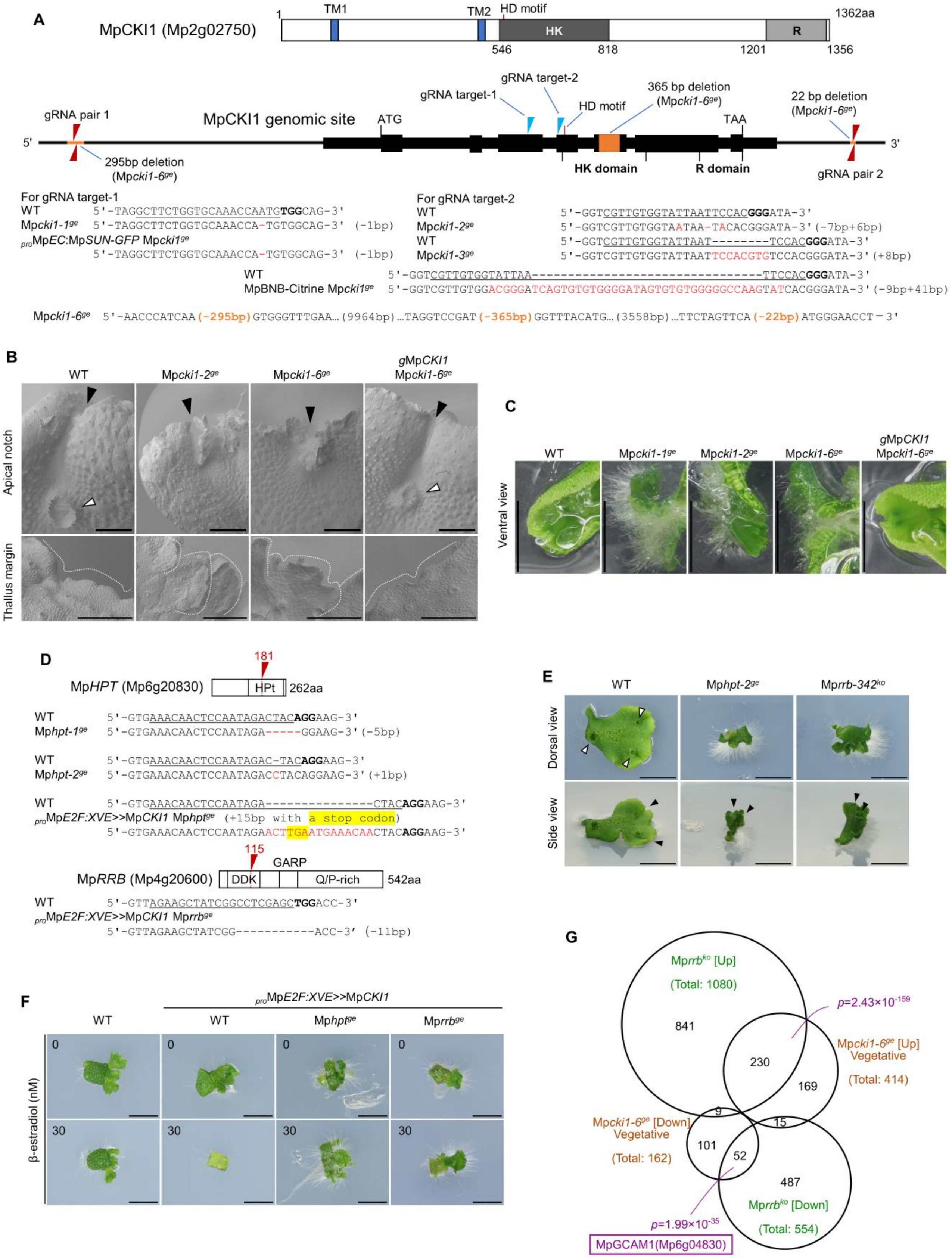
A
MpCKI1 (Mp2g02750)
1
TM1
TM2
HD motif
HK
R
1362aa
546
818
1201
1356
gRNA target-1
gRNA target-2
365 bp deletion
(Mpcki1-6ge)
22 bp deletion
(Mpcki1-6ge)
gRNA pair 1
MpCKI1 genomic site
ATG
HD motif
TAA
5'
3'
295bp deletion
(Mpcki1-6ge)
HK domain
R domain
gRNA pair 2
For gRNA target-1
WT 5'-TAGGCTTCTGGTGCAAACCAATGTGGCAG-3'
Mpcki1-1ge 5'-TAGGCTTCTGGTGCAAACCA-TGTGGCAG-3' (-1bp)
proMpEC:MpSUN-GFP Mpcki1ge
5'-TAGGCTTCTGGTGCAAACCA-TGTGGCAG-3' (-1bp)
For gRNA target-2
WT 5'-GGTCGTTGTGGTATTAATTCCACGGGATA-3'
Mpcki1-2ge 5'-GGTCGTTGTGGTAATAA-TACACGGGATA-3' (-7bp+6bp)
WT 5'-GGTCGTTGTGGTATTAAT--------TCCACGGGATA-3'
Mpcki1-3ge 5'-GGTCGTTGTGGTATTAATTCCACGTGTCCACGGGATA-3' (+8bp)
WT 5'-GGTCGTTGTGGTATTAA---------------------------------TTCCACGGGATA-3'
MpBNB-Citrine Mpcki1ge 5'-GGTCGTTGTGGACGGGATCAGTGTGTGGGGATAGTGTGTGGGGGCCAAGTATCACGGGATA-3' (-9bp+41bp)
Mpcki1-6ge 5'-AACCCATCAA (-295bp) GTGGGTTTGAA… (9964bp) …TAGGTCCGAT (-365bp) GGTTTACATG… (3558bp) …TTCTAGTTCA (-22bp) ATGGGAACCT-3'
B
WT
Mpcki1-2ge
Mpcki1-6ge
gMpCKI1
Mpcki1-6ge
Apical notch
Thallus margin
C
WT
Mpcki1-1ge
Mpcki1-2ge
Mpcki1-6ge
gMpCKI1
Mpcki1-6ge
Ventral view
D
181
MpHPT (Mp6g20830)
HPt
262aa
WT 5'-GTGAAACAACTCCAATAGACTACAGGAAG-3'
Mphpt-1ge 5'-GTGAAACAACTCCAATAGA-----GGAAG-3' (-5bp)
WT 5'-GTGAAACAACTCCAATAGAC-TACAGGAAG-3'
Mphpt-2ge 5'-GTGAAACAACTCCAATAGACCTACAGGAAG-3' (+1bp)
WT 5'-GTGAAACAACTCCAATAGA---------------CTACAGGAAG-3'
proMpE2F:XVE>>MpCKI1 Mphptge (+15bp with a stop codon)
5'-GTGAAACAACTCCAATAGAACTTGAATGAAACAACTACAGGAAG-3'
115
GARP
MpRRB (Mp4g20600)
DDK
Q/P-rich
542aa
WT 5'-GTTAGAAGCTATCGGCCTCGAGCTGGACC-3'
proMpE2F:XVE>>MpCKI1 Mprrbge
5'-GTTAGAAGCTATCGG-----------ACC-3' (-11bp)
E
WT
Mphpt-2ge
Mprrb-342ko
Dorsal view
Side view
F
proMpE2F:XVE>>MpCKI1
WT
WT
Mphptge
Mprrbge
0
30
β-estradiol (nM)
G
Mprrbko [Up]
(Total: 1080)
841
p=2.43×10^-159
230
Mpcki1-6ge [Up]
Vegetative
(Total: 414)
169
9
15
Mpcki1-6ge [Down]
Vegetative
(Total: 162)
101
52
487
Mprrbko [Down]
(Total: 554)
p=1.99×10^-35
MpGCAM1(Mp6g04830)

**Figure S2. Genotype information and vegetative phenotypes of mutants related to Mp*CKI1*, Mp*HPT*, and Mp*RRB* (related to Figure 1)**

A. Structure of MpCKI1 protein and genotype information for Mp*cki1$^{ge}$* mutants. TM, transmembrane domain. HK, histidine kinase domain; R, receiver domain; aa, amino acids. Blue arrowheads indicate two independent target sites for sgRNAs, and red arrowheads indicate target sites for gRNA pairs to generate Mp*cki1-6$^{ge}$*. The deleted regions in Mp*cki1-6$^{ge}$* are colored in orange.

B. SEM images of apical notches and margins of WT and Mp*cki1* thalli. Black arrowheads indicate apical notches, and open arrowheads indicate gemma cups. Dotted lines indicate thallus margins. Scale bars, 1mm (apical notch); 500 µm (thallus margin).

C. Ventral view of thalli grown on transparent gellan gum medium for rhizoid observation. Scale bars, 1 cm.

D. Structure of MpHPT and MpRRB and genotype information for gene-editing mutants. HPt, histidine-containing phosphotransfer domain; DDK, three phosphorylation sites in the signal-receiver domain of MpRRB; GARP, GARP domain, Q/P-rich, glutamine- and proline-rich domain. Red arrowheads indicate sgRNA target sites.

E. Dorsal and high-angle side view of 14-day-old thalli grown from explants containing apical notches, with genotypes as indicated. Black arrowheads indicate apical notches. Scale bars, 1 cm.

F. Regeneration assay showing 7-day-old thalli grown from explants without an apical notch, with indicated genotypes, of MpCKI1inducible overexpression lines ($_{\text{pro}}$*E2F:XVE*>>Mp*CKI1cds*) in the presence and absence of β-estradiol. Scale bars, 5 mm.

G. Venn diagram showing the comparison between differentially expressed genes (DEGs) in the thalli of Mp*cki1-6$^{ge}$* and Mp*rrb-307$^{ko}$*, each compared with the corresponding WT samples. Transcriptome data of Mp*rrb-307$^{ko}$* and the corresponding WT sample were retrieved from a previous study. [1] The transcriptomes were analyzed with the R package DESeq2, and DEGs were defined by *p*-values from a Wald test with Benjamini–Hochberg adjustment ($p$<0.01) and fold change in gene expression (|log2(Fold Change)|>0.585). Fisher's exact test was used to test if the overlap between two gene sets was significant.

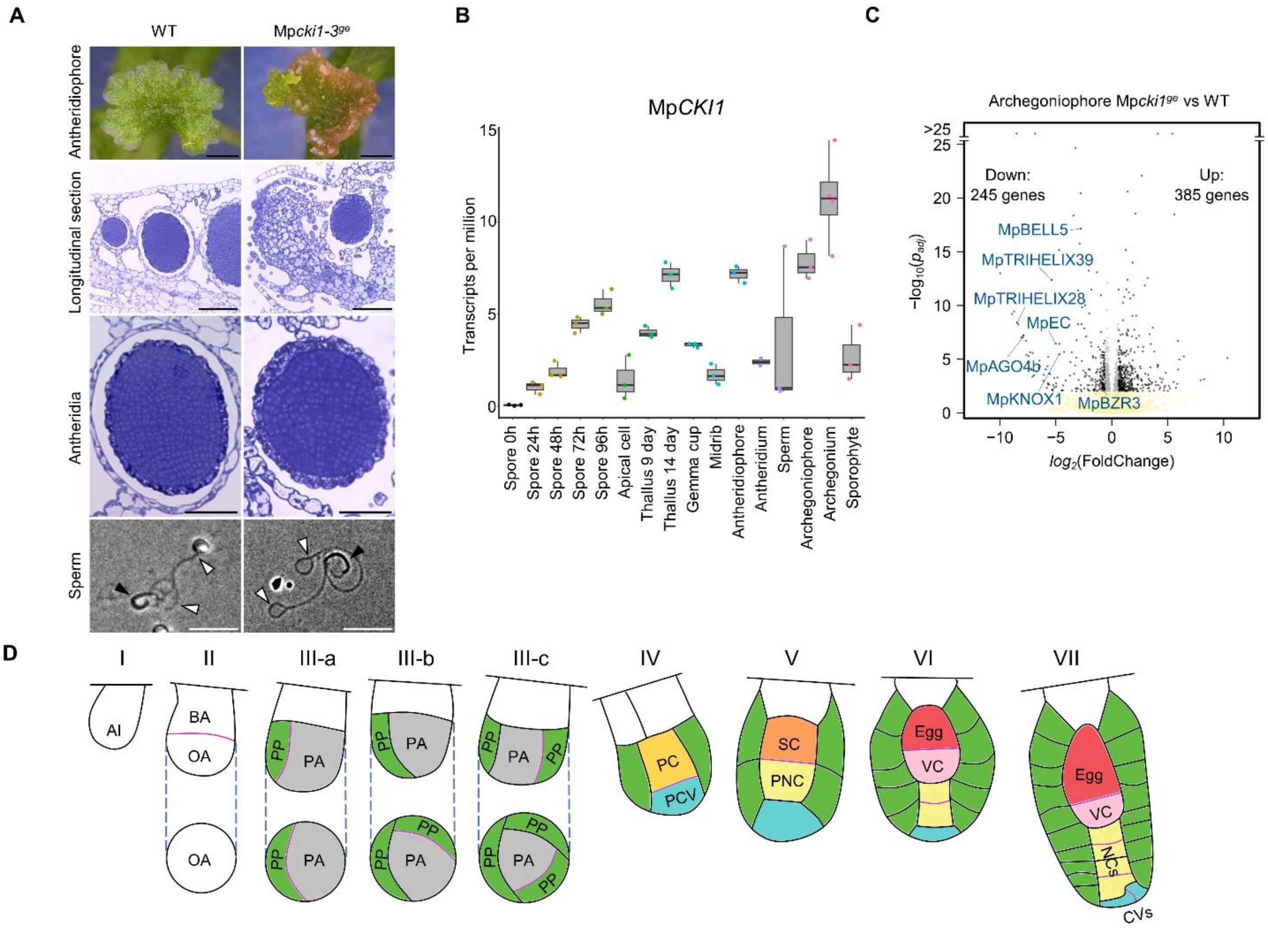
A
WT
Mpcki1-3ge
Antheridiophore
Longitudinal section
Antheridia
Sperm
B
MpCKI1
Transcripts per million
15
10
5
0
Spore 0h
Spore 24h
Spore 48h
Spore 72h
Spore 96h
Apical cell
Thallus 9 day
Thallus 14 day
Gemma cup
Midrib
Antheridiophore
Antheridium
Sperm
Archegoniophore
Archegonium
Sporophyte
C
Archegoniophore Mpcki1ge vs WT
Down:
245 genes
Up:
385 genes
MpBELL5
MpTRIHELIX39
MpTRIHELIX28
MpEC
MpAGO4b
MpKNOX1
MpBZR3
−log10(padj)
log2(FoldChange)
>25
D
I
II
III-a
III-b
III-c
IV
V
VI
VII
AI
BA
OA
PP
PA
PC
PCV
SC
PNC
Egg
VC
NCs
CVs

**Figure S3. Male sexual reproductive development of Mp*cki1* mutants, expression profiles of MpCKI1, archegoniophore transcriptome profiles of Mpcki*1* mutants, and nomenclature of cells in early archegonium development (related to Figure 2 and Figure 3)**

A. Male gametogenesis of WT and the Mp*cki1* mutant. Exposed antheridia are visible on the dorsal surface of the antheridiophore in the Mp*cki1* mutant. Open arrowheads indicate flagella of sperm, and black arrowheads indicate the main body of sperm. Scale bars, 2 mm (antheridiophore); 50 µm (longitudinal section and antheridia); 10 µm (sperm).

B. Expression profiles of Mp*CKI1* in different *M. polymorpha* tissues, summarized by MBEX (https://marchantia.info/mbex/) from previous studies. [2–9]

C. Volcano plot showing genes differentially expressed between Mp*cki1-6^ge^* and WT in archegoniophores. Points representing different genes were colored according to adjusted *p*-values by a Wald test (yellow if $-log_{10}(p_{adj})<1$) and fold change in gene expression (black if $|log_2(\text{Fold Change})|>0.585$, otherwise grey). Genes of interest are highlighted in blue and labeled with annotation.

D. Schematic illustration of archegonium development based on previous histological studies and our own observations. [10–12] Nomenclature is from Shimamura. [13] Coloring indicates different cells or cell lineages: green indicates the peripheral cell (jacket cell) lineage; yellow indicates the neck canal cell lineage; and blue indicates the cover cell lineage.

Stage I, a single Archegonial initial cell (AI) protrudes from the dorsal derivatives of the meristematic region of archegoniophore between the finger-like structures;

Stage II, AI divides transversely, results in a basal archegonial initial (BA, for the archegonial stalk and collar) and an outer archegonial initial (OA, for the archegonial venter and neck);

Stage III, OA undergoes three unequal vertical divisions and produces one primary axial cell (PA) surrounded by three primary peripheral cells (PPs, also named primary jacket cell in some studies);

Stage IV, PA is then unequally divided by a transverse wall to form a smaller primary cover cell (PCV) and a larger primary central cell (PC);

Stage V, PC divides transversely to form a secondary central cell (SC) and a PNC (primary neck canal cell);

Stage VI and VII, SC undergoes a transverse division to form an egg cell and a ventral canal cell (VC) in the venter. Meanwhile, the PNC successively divides via transverse walls to produce neck canal cells (NCs), and the PCV vertically divides to generate cover cells (CVs).

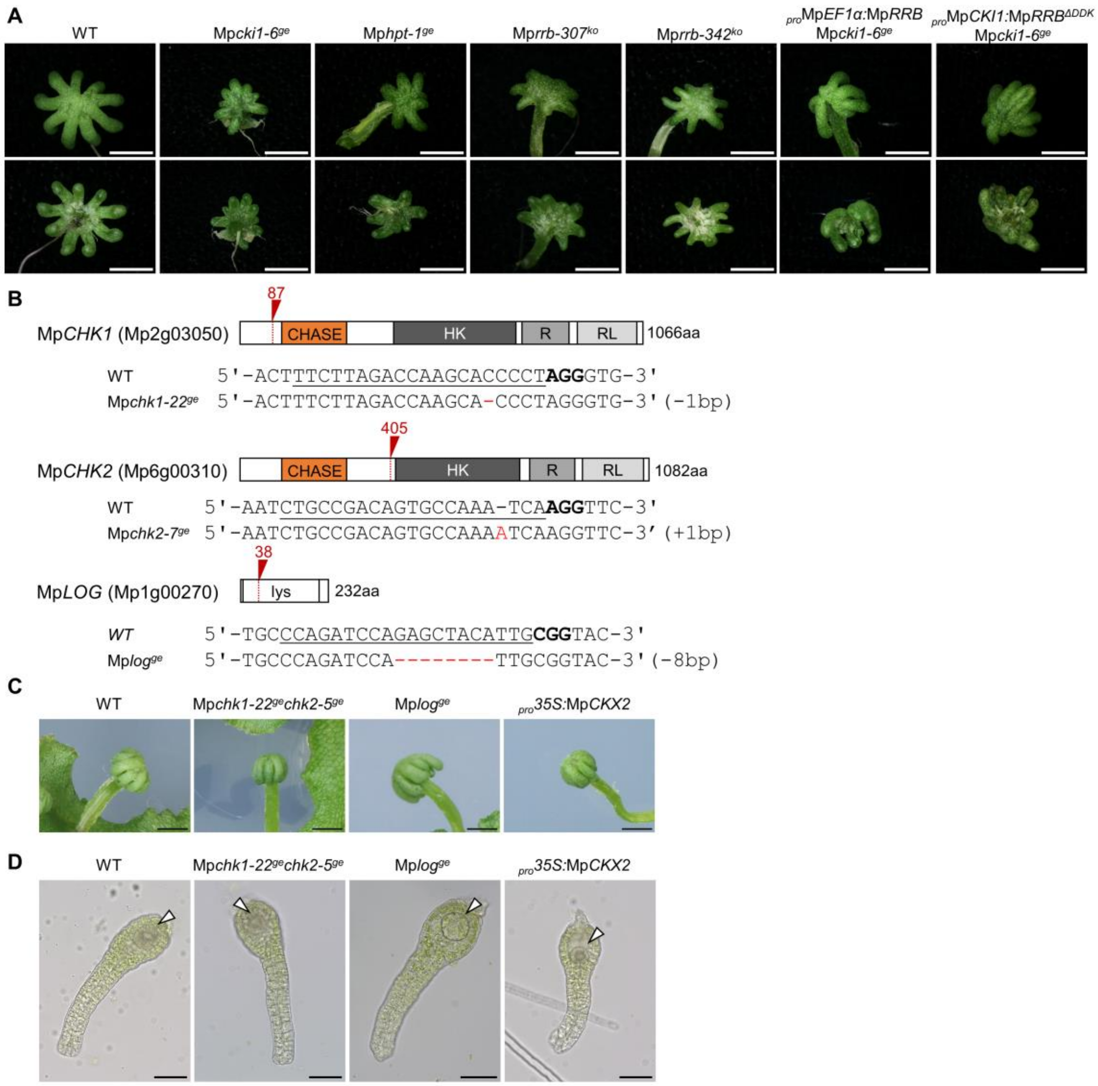


**Figure S4. Reproductive phenotypes related to TCS components and genotype information of mutants related to cytokinin signaling (related to Figure 4)**

A. Archegoniophore morphology related to the indicated genotypes. Scale bars, 2 mm.

B. Genotype information for double mutants of cytokinin receptors MpCHK1 and MpCHK2, and the mutation disrupting the gene encoding the cytokinin activation enzyme MpLOG. CHASE, Cyclases/Histidine kinases Associated Sensory Extracellular domain; HK, histidine kinase domain; R, receiver domain; RL, receiver-like domain; lys, lysine decarboxylase domain; aa, amino acids. Red arrowheads and dotted lines indicate sgRNA target sites.

C. Archegoniophore morphology in lines with the indicated genotypes, related to cytokinin reception and metabolism. Scale bars, 2 mm.

D. Isolated archegonia from lines with the indicated genotypes, related to cytokinin reception and metabolism. Empty arrowheads indicate egg cells. Scale bars, 50 μm.

**Table S3 List of oligonucleotides used in this study**

| Oligo name | Sequence (5' to 3') |
|---|---|
| **For gene editing (Mp*cki1*)** | |
| MpCKI1-sgRNA#L-F | CTCGGCTTCTGGTGCAAACCAATG |
| MpCKI1-sgRNA#L-R | AAACCATTGGTTTGCACCAGAAGC |
| MpCKI1-sgRNA#R-F | CTCGCGTTGTGGTATTAATTCCAC |
| MpCKI1-sgRNA#R-F | AAACGTGGAATTAATACCACAACG |
| MpCKI1-ld-sgRNA1-F | TGCGCTGAACTTAGCAGCAA |
| MpCKI1-ld-sgRNA1-R | AAACTTGCTGCTAAGTTCAGCGCA |
| MpCKI1-ld-sgRNA2-F | CTCGGGATGAACTAGAACCTTTCC |
| MpCKI1-ld-sgRNA2-R | AAACGGAAAGGTTCTAGTTCATCC |
| MpCKI1-ld-sgRNA3-F | CTCGCAAGCCGGAACTCCTCGGTT |
| MpCKI1-ld-sgRNA3-R | AAACAACCGAGGAGTTCCGGCTTG |
| MpCKI1-ld-sgRNA4-F | CTCGCCTTTAAACCTGTTATTTAT |
| MpCKI1-ld-sgRNA4-R | AAACATAAATAACAGGTTTAAAGG |
| **For *g*Mp*CKI1* cloning** | |
| MpCKI1_5'_EcoRI_IFF | ATCCGGTACCGAATTCCGCCTGGTCATGTGAATCTT |
| MpCKI1_3'_EcoRI_IFR | GTGCGGCCGCGAATTCTTTCCGATCTTTCTCGAGGAG |
| **For Mp*CKI1* promoter cloning** | |
| MpCKI1_pro_FB4 | GGGGACAACTTTGTATAGAAAAGTTGCGCCTGGTCATGTGAATCTT |
| MpCKI1_pro_RB1R | GGGGACTGCTTTTTTGTACAAACTTGCTCTCCTATACTCTTCCCACA |
| **For Mp*CKI1* CDS cloning** | |
| MpCKI1-CDS-F | CACCATGGAAGGGCGGGGA |
| MpCKI1-CDS-R | TTAACTAGGTCGGTTGCCTG |
| **For gene editing (Mp*hpt*)** | |
| MpHPT-sgRNA-F | CTCGAAACAACTCCAATAGACTAC |
| MpHPT-sgRNA-R | AAACGTAGTCTATTGGAGTTGTTT |
| **For gene editing (Mp*rrb*)** | |
| MpRRB-sgRNA-F | CTCGAGAAGCTATCGGCCTCGAGC |
| MpRRB-sgRNA-R | AAACGCTCGAGGCCGATAGCTTCT |
| **For Mp*RRBcds*$^{ΔDDK}$ cloning** | |
| MpRRBΔDDK-CACCATG-F | CACCATGGGACGAGGCGAAACGATGAA |
| MpRRB-nstop-R | CTGACCCTGACCAGGCCTAT |
| **For gene editing (Mp*chk1*&Mp*chk2*)** | |
| MpCHK1-sgRNA-F | CTCGTTCTTAGACCAAGCACCCCT |
| MpCHK1-sgRNA-R | AAACCCTAGGGGTGCTTGGTCTAAGAA |
| MpCHK2-sgRNA-F | CTCGCTGCCGACAGTGCCAAATCA |
| MpCHK2-sgRNA-R | AAACTGATTTGGCACTGTCGGCAG |
| **For gene editing (Mp*log*)** | |
| MpLOG-sgRNA-F | CTCGCCAGATCCAGAGCTACATTG |

| | |
|---|---|
| MpLOG-sgRNA-R | AAACCAATGTAGCTCTGGATCTGG |

Red letters indicate adapter sequences, and underlined sequences indicate start codons.
